# Revisiting tolerance to ocean acidification: insights from a new framework combining physiological and molecular tipping points of Pacific oyster

**DOI:** 10.1101/2021.09.21.461261

**Authors:** Mathieu Lutier, Carole Di Poi, Frédéric Gazeau, Alexis Appolis, Jérémy Le Luyer, Fabrice Pernet

## Abstract

Studies on the impact of ocean acidification on marine organisms involve exposing organisms to future acidification scenarios as projected for open ocean, which has limited relevance for coastal calcifiers. Characterization of reaction norms across a range of pH and identification of tipping points beyond which detrimental effects are observed has been limited and focus on only a few macro-physiological traits. Here we filled this knowledge gap by developing a framework to analyze the broad macro-physiological and molecular responses over a wide pH range of juvenile Pacific oyster, a model species for which the tolerance threshold to acidification remains unknown. We identify low tipping points for physiological traits at pH 7.3-6.9 that coincide with a major reshuffling in membrane lipids and transcriptome. In contrast, shell parameters exhibit effects with pH drop well before tipping points, likely impacting animal fitness. These findings were made possible by the development of an innovative methodology to synthesize and identify the main patterns of variations in large -omic datasets, fit them to pH and identify molecular tipping-points. We propose the application of our framework broadly to the assessment of effects of global change on other organisms.

## Introduction

The exponential increase in the atmospheric emission of carbon dioxide (CO_2_) from anthropogenic activities is mitigated by ocean absorption that leads to decreasing surface ocean pH and changes in carbonate chemistry, a phenomenon known as ocean acidification (OA) (Caldeira & Wickett, 2003; Orr et al., 2005). This represents a tremendous challenge for marine organisms, especially for calcifiers that produce calcium carbonate (CaCO_3_)-based exoskeletons. OA not only induces internal acidosis that impacts metabolism, behavior, growth and reproduction, but also decreases carbonate ions (CO_3_^2-^) concentration in the ocean, the elemental constituent of calcifier exoskeletons (Gazeau et al., 2013; IPCC, 2019; Kroeker et al., 2013; Tresguerres & Hamilton, 2017).

OA has been the most-studied topic in marine science in recent times (Browman, 2017). As such, many OA experiments have been conducted, usually exposing organisms to experimental conditions based on scenarios modelled for open ocean waters, typically simulating present and near-future surface ocean pH levels. However, many calcifying species thrive in coastal areas where pH levels vary far more than in the open ocean on both daily and seasonal scales (Vargas et al., 2017; Waldbusser & Salisbury, 2014). Such variability in coastal environments limits the relevance of applying projections based on simulations for the open ocean and indicates that knowing the physiological response of coastal organisms over a wider pH range (Dorey et al., 2013) would be necessary to accurately assess OA impact.

Reaction norm, i.e. the response of an organism to changing environmental parameters, allows the identification of tipping points that beyond small variations will have major impacts. According to the last IPCC reports, identification of tipping-points is a key knowledge gap in environmental change research (IPCC, 2019, 2021). To date, only four studies have experimentally established reaction norms for marine calcifiers in relation to pH. These studies focused on the assessment of tipping points for a few selected traits measured at the organism level (i.e. growth, survival) and provide a limited view of the whole organism response, ignoring the molecular level (Comeau et al., 2013; Dorey et al., 2013; Lee et al., 2019; Ventura et al., 2016). We therefore identify the need to integrate macro-physiology and omics together with the reaction norm for understanding the mechanisms responsible for individual organism success under a changing environment. This however requires developing an innovative methodology to synthesize and identify the main patterns of variations in large -omics datasets (Strader et al., 2020) and fit them to reaction norm.

Transcriptomic and lipidomic approaches provide holistic views of organismal responses to environmental changes at the molecular and biochemical levels. Organisms do vary in their transcriptomic responses to pH but common patterns have been observed, including shifts in acid-base ion regulation, metabolic processes, calcification and stress response mechanisms (Matz, 2018; Strader et al., 2020). Membrane composition in terms of fatty acid is also modulated as a response to environmental changes and influence exchanges between intra and extracellular compartments with impacts on metabolic rates (Hazel & Williams, 1990; Hochachka & Somero, 2002; Hulbert & Else, 1999).

Here we determine the reaction norm of juvenile Pacific oyster *Crassostrea gigas*, one of the most cultivated invertebrate species in the world (FAO, 2020), over a wide range of pH for macro-physiological traits, membrane lipids and gene expression. Although the impacts of OA on this model species have been intensively studied using scenario approaches, there is currently no consensus on its robustness to low pH (Ducker & Falkenberg, 2020). The establishment of reaction norm with biochemical and molecular data is a novel approach that provide new insights on the sensitivity of marine calcifiers to OA.

## Material and methods

### Animals and maintenance

The oysters were produced at the Ifremer hatchery facilities in Argenton (Brittany, France) in late August 2018 according to Petton et al. (2015). The broodstock consisted of 139 females and 40 males originating from six different cohorts collected in the natural environment between 2011 and 2016, off Fouras (Ile d’Aix, France). At 40 days old, the juveniles were moved to the Ifremer growing facilities in Bouin (Vendée, France). On January 10^th^ 2019, oysters were returned to Argenton and kept for 8 d in a 500 L flow-through tank. Seawater temperature was gradually increased from 7 °C to 22 °C, the optimal temperature for *C. gigas* (Bayne, 2017), at a rate of *ca*. 2 °C d^-1^. During the experiment, oysters were continuously supplied with natural seawater originating from a pool (∼9000 m^3^) which is renewed with each spring tide, filtered at 5 µm and UV-treated. The oysters were fed continuously on a mixed diet of *Isochrysis affinis galbana* (CCAP 927/14) and *Chaetoceros gracilis* (UTEX LB2658) (1:1 in dry weight). Food concentration was maintained at 1500 μm^3^ mL^−1^ of phytoplankton cells at the outlet of the tank – *ad libitum* (Rico-Villa et al., 2009) – and controlled twice daily using an electronic particle counter (Multisizer 3, Beckman Coulter, USA) equipped with a 100 μm aperture tube. On the eve of the experiment on January 14^th^ 2019, oysters were 5-month-old and divided into 15 batches containing 292 ± 20 individuals (95.2 ± 0.2 g).

### Experimental design

Each batch of oysters was exposed to one constant nominal pH condition ranging from pH 7.8 to 6.4 with a step of 0.1 between two levels. The upper pH condition (pH ∼7.8) was obtained by running seawater with oysters without pH regulation. The experimental system consisted of 18 experimental units that were randomly assigned to one pH condition (n = 15) or to control blanks without animals (n = 3). Each experimental units consisted of a header tank in which seawater was acidified (except for the ambient pH and blanks) and then delivered by a pump to a holding tank containing the oysters. These tanks were 45 L and their entire volume was renewed every 82 min. pH was regulated by means of pure-CO_2_ bubbling, controlled by a pH-regulator (ProFlora^®^ u403 JBL). The pH-regulator was connected to pH-probes installed in each header tank (pH-sensor+Cal, JBL), checked at the start of the experiment using Certipur^®^ Merck NBS buffers (pH = 4.00, pH = 7.00). pH electrodes from the pH-regulator were inter-calibrated twice daily against a pH probe (Sentix^®^ 940-3 WT) connected to a Multiline^®^ Multi 3630 IDS-WTW. The pH-probe was checked once a week using Certipur^®^ Merck NBS buffers (pH = 4.00, 7.00, and 9.00) and calibrated twice a week on the total scale with a certified Tris/HCl buffer (salinity 35; provided by A. Dickson, Scripps University, USA). Throughout the text, pH levels are therefore expressed on the total scale (pH_T_). In each experimental tank, air bubbling and a small homogenisation pump (3 W) ensured an efficient mixing of seawater surrounding the oysters. Photoperiod was 10 h light: 14 h dark. During the entire experimental period, oysters were fed as described above.

On January 18^th^ 2019, each batch of oyster was randomly assigned to one tank. Oysters were first held at ambient pH for 3 d. Then, pH was progressively decreased in each pH-regulated tank at a rate of 0.2 unit d^-1^. The decrease in pH lasted for 7 d for the lowest condition. Experimental pH conditions were all reached on January 27^th^ 2019 and oysters were maintained at these experimental pH levels for 23 d. No mortality was recorded during the experiment.

### Seawater carbonate chemistry

In each oyster tank, temperature, salinity, dissolved oxygen (O_2_) saturation levels and pH were measured twice a day using a Multiline^®^ Multi 3630 IDS-WTW (pH probe Sentix^®^ 940-3 WTW, O_2_ probe FDO^®^ 925 WTW, salinity probe TetraCon^®^ 925 WTW). Seawater samples were collected weekly, filtered on 0.7 μm (GF/F, Whatman^®^) and poisoned with 0.05% mercury (II) chloride. Total alkalinity (TA) was then measured in triplicate 50 mL subsamples by potentiometric titration at 22 °C, using a Titrando 888 (Metrohm^®^) titrator coupled to a glass electrode (ecotrode plus, Metrohm^®^) and a thermometer (pt1000, Metrohm^®^). TA was calculated following the protocol described in Dickson et al. (2007). pH and TA, together with salinity and temperature, were used to determine carbonate chemistry parameters using the package seacarb v 3.2.16. of the R software.

### Biometry

The total fresh weight (shell + tissue) of each batch of oysters was measured twice weekly and interpolated between measurements to estimate daily weight. In addition, shell length and total fresh weight were measured individually on 30 oysters from each condition at the onset (1 d) and at the end of the experiment (23 d). These oysters were then dissected and pooled to determine the total weight of shell and tissue, separately. The tissues were then lyophilized and weighted to obtain dry weight. Growth rate was calculated as:

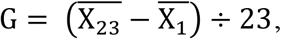

where *G* is growth rate as expressed as increase in shell length or total body weight per day (mm d^-1^, g d^-1^), 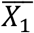and, 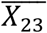 are the average parameter values for shell length and total weight measured at the onset (1 d) and the end of the experiment (23 d).

Shell thickness was measured on the left valve of five individuals collected per pH condition at the end of the experiment. The left valve was more susceptible to acidification than the right valve. We observed holes in the shell under low pH condition (pH < 6.6) that always occurred on the left valves, close to the umbo. Shells were dried for 24 h at 45 °C, embedded in polyester resin and cross-sectioned from umbo to opposite shell margin along the longitudinal growth axis using a precision saw (Secotom-10 Struers). Sections were glued on a microscope slide and polished with silicon carbide abrasive disks (1200 and 2500 grains cm^-2^). Images of the section were captured under a Lumar V12 stereoscope (Zeiss) at 30x magnification, and the entire section was reconstructed using an image acquisition software (AxioVision SE 64 – v4.9.1, Zeiss). The minimal shell thickness was measured within the first third of the shell starting from umbo using ImageJ software (Schneider et al., 2012).

### Physiological rates

Seawater was sampled twice a day at the inlet and outlet of each oyster tank and phytoplankton cell concentrations were measured using an electronic particle counter (see previous section). No pseudofaeces production was detected throughout the experiment. Ingestion rate was determined as:

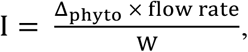

where *I* is the ingestion rate expressed as cm^3^ min^-1^ g^-1^, *Δ*_*phyto*_ is difference in phytoplankton concentrations between the inlet and the outlet of the oyster tanks (cm^3^ min^-1^), *flow rate* is the water flow at the inlet of the oyster tanks (mL min^-1^), and *W* is the total fresh weight of oyster batch (g). The three control blanks were used to check that there was no sedimentation of algae (no differences in cell concentrations between the inlet and the outlet, data not shown).

Net calcification and respiration rates were measured after 22 d of exposure. Food supply was stopped 17 h before the assay and tanks were emptied, cleaned and refilled. Again, 1 h before the assay, the tanks were emptied, cleaned and refilled. At the onset of the incubation, water flow and air bubbling were stopped. Gentle mixing of the seawater was maintained by homogenization pump but air bubbling was stopped. Incubations lasted for 90 min. This duration allowed to keep the pH close to the setpoint (< 0.1 pH unit variation) despite oyster respiration. At the onset and at the end of the incubation period, temperature, salinity, pH and O_2_ concentration (mg L^-1^) were measured, and seawater samples were filtered and poisoned for TA analyses (see above) or were immediately frozen at -20 °C for ammonium (NH_4_^+^) measurements (Aminot & Kérouel, 2007) (see Supplementary T154:161able 4). Empty tanks were used as controls to check that there was no change in any of the parameters due to evaporation or other potential biological processes in the water itself. Net calcification rate was determined following a modified procedure from Gazeau et al. (2015) using the alkalinity anomaly technique (Smith & Key, 1975):

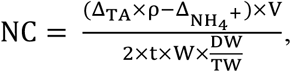

where *NC* is the net calcification rate expressed as µmol CaCO_3_ h^-1^ g^-1^, *Δ*_*TA*_ and *Δ*_*NH4+*_ are differences in TA (*µ*mol Kg^-1^) and NH_4_^+^ (µmol L^-1^) between the onset and the end of the incubation period, *ρ* is seawater density (kg L^-1^) calculated based on temperature and salinity during the incubation, *V* is the volume of the tanks (L), *t* is the duration of the incubations (h), 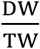 is the ratio of dry weight and total weight determined after lyophilization of a pool of tissues from 30 oysters (see above). Respiration rate was determined following:

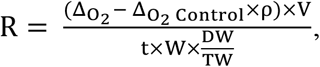

where *R* is the respiration rate expressed as mg O_2_ h^-1^ g^-1^, 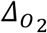 and 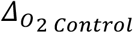 are differences in O_2_ concentration (mg L^-1^) between the onset and the end of the incubation period in the oyster tank and in the three control tanks (average) respectively.

### Biochemistry

Soft-tissues of five individuals from each pH condition were collected at 23 d, flash-frozen in liquid nitrogen, pooled, grounded with a ball mill and stored at -80 °C pending analyses. Oyster powder was diluted with chloroform/methanol (2:1, v/v) for the determination of neutral lipids (triacylglycerol: TAG, and sterols: ST) using high performance thin layer chromatography. TAG-ST ratio was used as a proxy of the relative contribution of lipid reserve to structure (membrane). Polar lipids were purified on silica gel micro column, transesterified using methanolic H_2_SO_4_ at 100 °C, and the resulting fatty acid methyl esters (FAME) were analyzed using a gas-chromatography flame ionization detection system equipped with a DB-Wax capillary column. Peaks were analyzed by comparison with external standards. Each fatty acid was expressed as the peak area percentage of total polar fatty acids. Carbohydrate content (μg mg^−1^) was determined according to the colorimetric method described in DuBois et al. (1956).

### Transcriptomics

Soft-tissues of five individuals from each tank were collected at 23 d, flash-frozen in liquid nitrogen and individually grounded (n = 75). Total RNA was then extracted with Extract-All (Eurobio) at a ratio of 5 mL per 100 mg of tissue. RNA quantity/integrity and purity were verified with a NanoDrop 2000 spectrophotometer (Thermoscientific®) and a Bioanalyzer 2100 (Agilent Technologies®) respectively. Samples were then DNase-treated using a DNase Max™ Kit (MO BIO Laboratories, Inc.) and analyzed at the Genotoul sequencing platform (INRAE US 1426 GeT-PlaGe, Centre INRAE de Toulouse Occitanie, Castanet-Tolosan, France). TruSeq RNA libraries were multiplexed and sequenced on a single lane of NovaSeq6000 Illumina S4 150-bp paired-end.

Raw reads were first filtered using Trimmomatic v.0.36 for a minimum length (60 bp), a minimum quality (trailing = 20, leading = 20) and the presence of potential contaminants and remaining adaptors (https://ftp.ncbi.nlm.nih.gov/pub/UniVec; 08/17/20). The quality of the reads was monitored before and after this trimming process with FastQC v.0.11.5 (https://www.bioinformatics.babraham.ac.uk/projects/fastqc/). The *C. gigas* reference genome (Peñaloza et al., 2021) was downloaded from NCBI (GCF_902806645.1) and indexed with Gmap v2020.06.01. Filtered reads were mapped against the reference genome using GSNAP v2020-06-30 keeping default parameters but allowing a minimum mismatch value of 2 and a minimum read coverage of 90%. Finally, the gene expression levels were assessed using HTSEQ v0. 6.1. The DESeq2 v1.30.0 R package was used to process the expression with a first step of normalization using the variance-stabilizing transformation implemented in the *‘vst’* function. The *vst* matrix was used to build a signed weighted co-expression network analysis implemented in the WGCNA v1.69 R package. First, genes with low overall variance (< 5%) were removed for the analysis as recommended (Langfelder & Horvath, 2008). We fixed the “soft” threshold powers of 13 the scale-free topology criterion to reach a model fit (|R|) of 0.90. Clusters were defined using the “*cutreeDynamic*” function (minimum of 50 genes by cluster and default cutting height of 0.99) based on the topological overlap matrix, and an automatic merging step with the threshold fixed at 0.25 (default) allowed merging correlated clusters. For each cluster, we defined the cluster membership (kME; Eigengene-based connectivity) and only clusters showing significant correlation (p < 0.01) to pH were kept for downstream functional analysis.

### Statistical analysis

All statistical analyses were performed using the R software v4.0.3 and the threshold of statistical significance was fixed at 0.05 unless specifically stated. Relationships between dependent variables and average pH recorded during the exposure period (or the incubation period for net calcification and respiration rates) were computed using regression models. Dependent variables were biometrics (shell length, total body weight, and growth), physiological rates (ingestion, net calcification and respiration), biochemistry (lipid reserves, carbohydrates) and gene expression. Each gene included in the WGCNA clusters were individually tested against pH. Fatty acids were summarized using principal component analysis and separated into two groups according to their correlation with first principal component (positive or negative). Each group was then summed and tested against pH. Piecewise, linear and log-linear regression models were tested and compared for each individual variable (or set of variables). In each case, the model that had the highest R^2^, the lowest Akaike and Bayesian information criteria (AIC, BIC) and the lowest residual sum of squares was selected. For each piecewise regression model, we estimated the tipping point, defined as the values of the pH where the dependent variables tip, by implementing the bootstrap restarting algorithm (Wood, 2001). For physiological rates, we also determined the critical-point defined as the pH at which the dependent variable was equal to zero. Normality of residual distribution was tested using Shapiro-Wilk test and homogeneity of variances was graphically checked. Significance of each slope was tested according to Student’s *t* test. Piecewise regression models were built using the segmented v1.3-4 R package.

For transcriptomic analysis, only genes that are significantly correlated to pH were tested among the retained clusters (Pearson correlation). The frequency distribution of pH tipping-point values was plotted for each cluster of genes. Groups of genes which exhibit neighboring tipping points with distribution frequencies > 5%, were grouped together and used for GO enrichment analysis using GOA-tools v0.7.11 (Klopfenstein et al., 2018), implemented in the Github repository ‘*go_enrichment’* (https://github.com/enormandeau/go_enrichment) and the go-basic.obo database (release 2019-03-19) with Fisher’s extract tests. Only significant GO terms that included a minimum of three genes and with Bonferroni adjusted P < 0.01 were kept. For complementarity with previous studies, we also specifically examined the genes that are commonly reported as involved in the calcification process and in the formation of the organic matrix of the shell and periostracum.

## Results

### Acclimation of oysters to 15 different pH conditions

During the experiment, pH levels in the oyster tanks were stable and reached the targeted values, except for the lowest pH condition that was 6.5 ± 0.3 instead of 6.4 (Supplementary Note 1, Supplementary Table 1, Supplementary Figure 1). The effects of acidification were clearly visible at the end of the experiment on the color and the size of the shells, as shown in Figure 1. Shell pigmentation apparently decreased with decreasing pH. Total body weight and shell length of oysters ranged from 0.4 ± 0.1 to 2.0 ± 0.8 g and from 12.9 ± 2.1 to 25.4 ± 4.5 mm respectively (mean ± SD; Figure 2A, Supplementary Figure 2).

**Figure 1.**
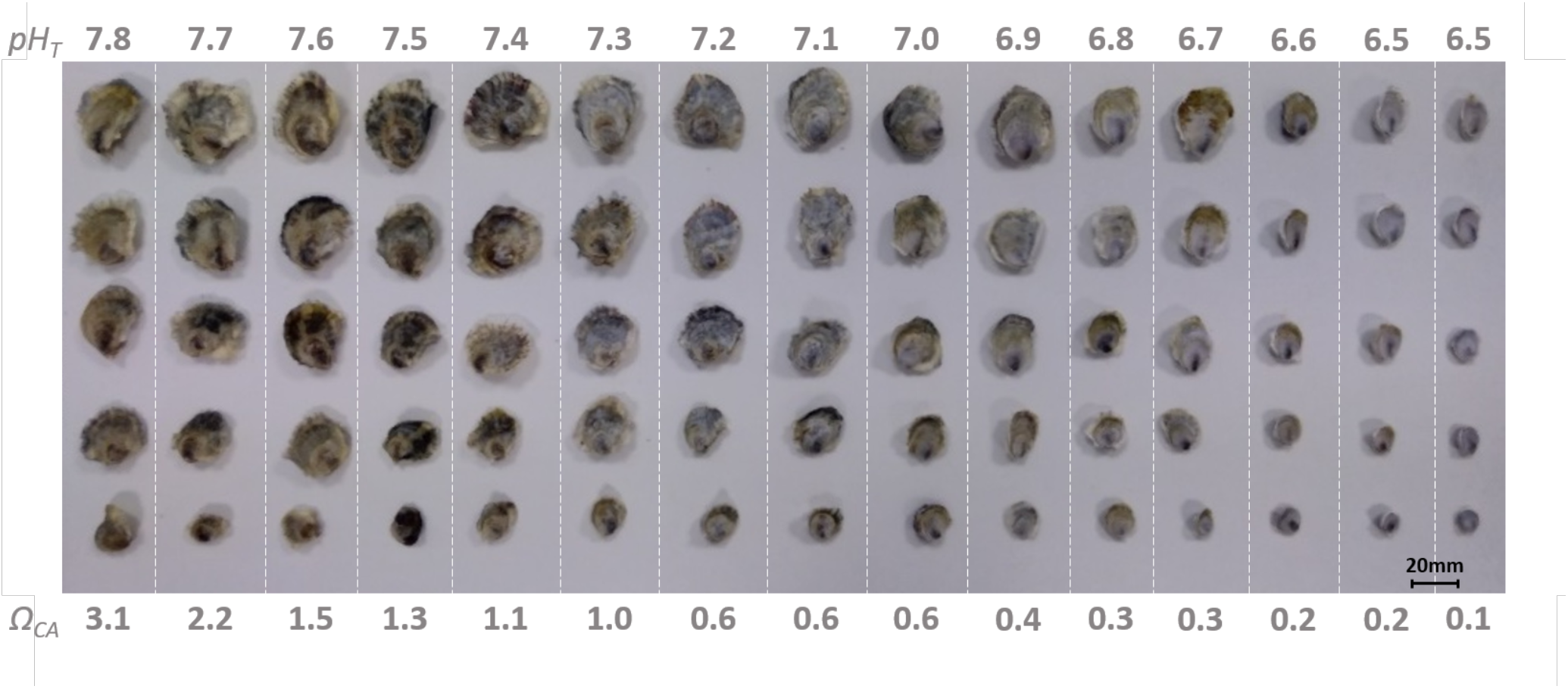
Oysters exposed to 15 pH conditions for 23 d. Five oysters were selected from each condition, and sorted from the smallest to the biggest. Corresponding pH (total scale) and saturation states of seawater with respect to calcite (Ω_CA_) are shown.

**Figure 2.**
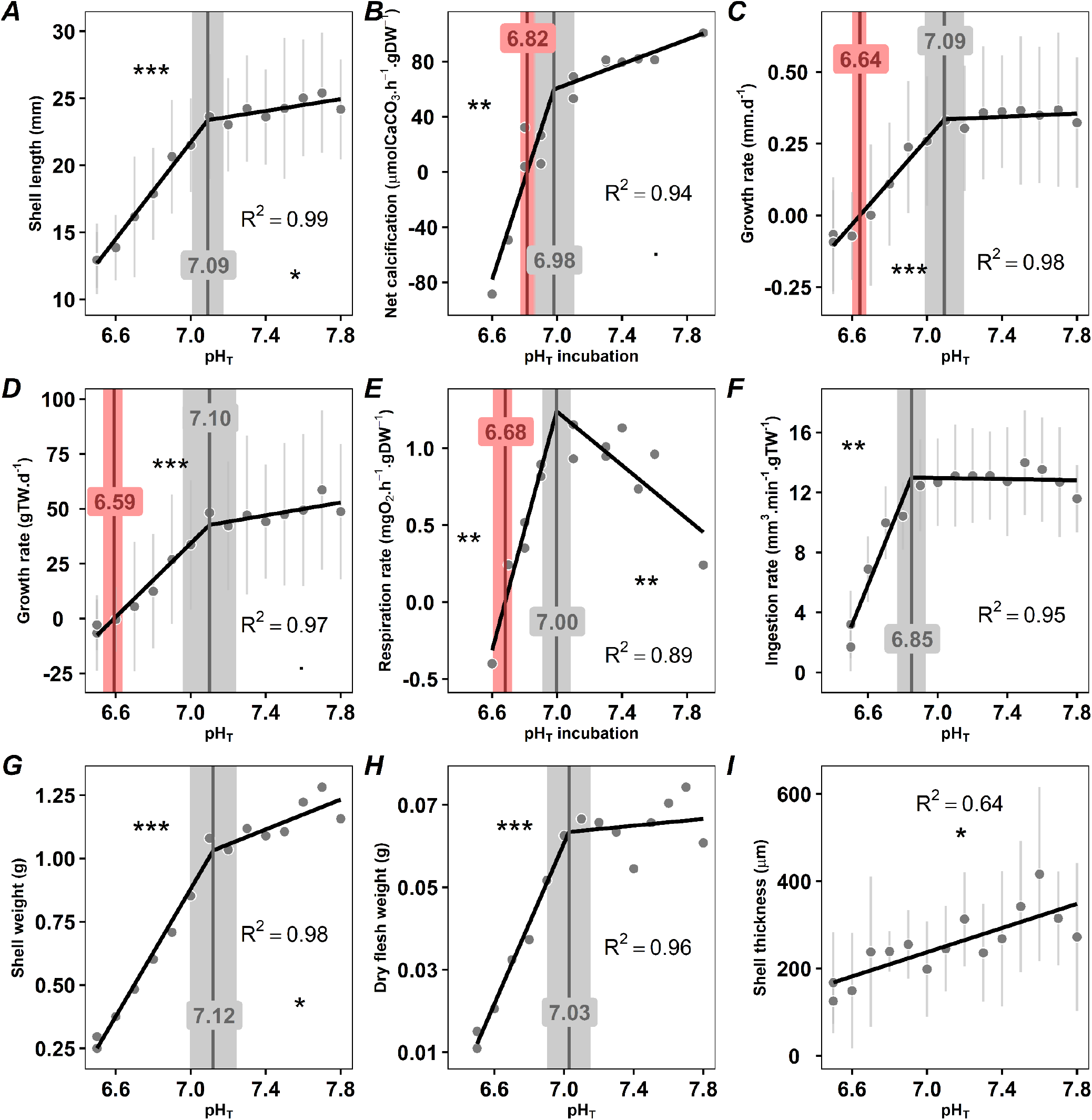
Biometry and physiological rates of oysters as a function of pH (total scale). (A) shell length, (B) shell thickness, (C) shell weight, (D) growth rate in shell length, (E) growth rate in total body weight (TW), (F) dry flesh weight, (E) net calcification rate, (H) ingestion rate and (I) respiration rate. Data are means ± SD when available. Tipping points and critical points and their 95% confidence intervals are shown in grey and red, respectively. The significance levels of the slopes are presented using symbols (P< 0.001 ***, < 0.01 **, < 0.05 *, < 0.1 .).

### Most macro-physiological traits tip at low pH values between 7.1-6.9

We measured biometric parameters like the length, thickness, and weight of the shell and the total body weight and the dry flesh weight at the end of the 23-day exposure to the 15 different pH conditions. We calculated growth rates for shell length and total body weight, and we measured physiological rates like calcification, ingestion and respiration rates. From these data, we identified an overall tipping point for macro-physiological traits at pH ∼7.1-6.9 below which they declined sharply (Figures 2A-H). Calcification rate and growth rates exhibited critical points at pH 6.8 and 6.6, respectively, below which they became negative and turned to net dissolution of the shell and weight losses (Figures 2B-D). Concomitantly, respiration rate was arrested at pH 6.7 and ingestion rate was near-zero at pH 6.5 (Figure 2E-F).

### Shell parameters and respiration rate are impacted above tipping point

Although macro-physiological traits were generally unaffected by reductions in pH above tipping points, this was not the case for shell length, shell weight and respiration rate. Both shell parameters indeed decreased with decreasing pH while the respiration rate increased before reaching the tipping point (Figures 2A, 2E and 2G). In addition, the shell thickness was the only parameter that decreased linearly over the entire pH range without tipping point (Figure 2I).

### A major remodeling of membrane lipids occurs at pH 6.9

To provide an in-depth characterization of the reaction norms at the whole organism level, we aimed to link the oyster reaction norm established for the macro-physiological traits described previously to the molecular responses of oysters. We first analyzed the fatty acid composition of membrane lipids that play an important role in exchanges between intracellular and extracellular compartments. Principal component analysis of all fatty acids showed that the first axis (PC1) alone explained 66% of the total variance in relation to pH (Figure 3A). The set of fatty acids was reduced to two terms defined by the sums of fatty acids that correlated with PC1 either positively or negatively, and each term was plotted against pH (Figure 3B-C). Positively correlated fatty acids mainly consisted of docosahexaenoic acid (DHA, 22:6n-3) and palmitic acid (PA, 16:0) contributing 43% and 14%, respectively, to PC1 (Supplementary Table 2). This group of fatty acids exhibited a tipping point at pH 6.9, below which their contribution to membrane decreased markedly (Figure 3B) at the benefit of the fatty acids that were negatively correlated with PC1. These negatively correlated fatty acids consisted of arachidonic acid (ARA, 20:4n-6), eicosapentaenoic acid (EPA, 20:5n-3) and non-methylene interrupted fatty acids (22:2NMI_i,j_), which contributed 12%, 6% and 6%, respectively, to PC1 (Figure 3C) (Supplementary Table 2).

**Figure 3.**
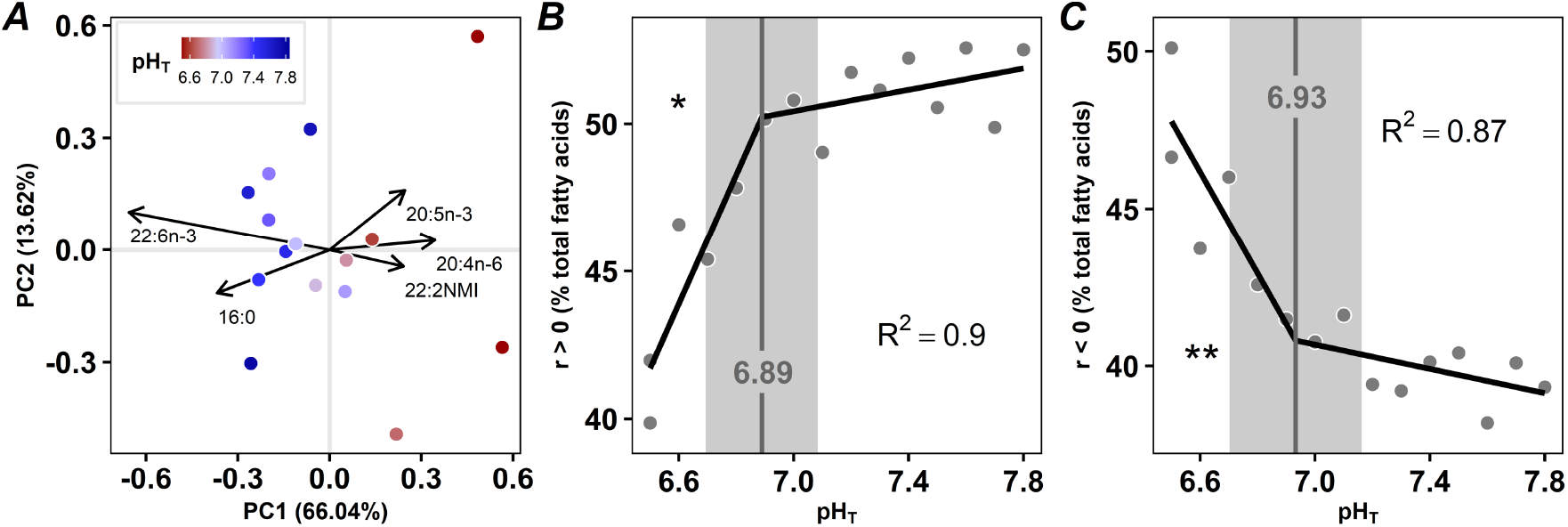
Membrane fatty acid (FA) composition of oysters as a function of pH (total scale). (A) Principal component analysis of polar fatty acid classes for oysters (n = 15 pools of five oysters) exposed to 15 pH levels. Arrows represent fatty acids contributing to more than 5% of the first principal component (PC1). Individuals are colored according to the average pH. Contribution to membrane of the sum of fatty acids that are (B) positively or (C) negatively correlated to PC1 as a function of pH. Tipping points and their 95% confidence intervals are shown in grey. The significance levels of the slopes are presented using symbols (P < 0.001 ***, < 0.01 **, < 0.05 *, < 0.1 .).

### Expression of most genes shows tipping points at pH 7.3-6.9

To further assess the molecular effects of pH change, we compared by RNA-seq the transcriptomic responses of oysters exposed to the 15 pH conditions for 23 days. To do this, we first clustered the differentially expressed genes that co-vary together using WGCNA analysis and retained the clusters that were correlated with pH (Supplementary Table 3). Then, we plotted the frequency distribution of pH tipping-point for each cluster of genes (Figure 4). Owing to this original method, we found that 1054 genes, distributed in three clusters, showed linear (or loglinear) or piecewise regression with pH (Supplementary table 3). Among them, 49% showed tipping-points at pH 7.3-6.9 (Figure 4A-C). Expression level of most genes were unchanged between pH conditions down to the pH tipping point, below which it increased for genes in clusters 1 and 2 (Figure 4A-B), or decreased for genes in cluster 3 (Figure 4C). Some genes linearly increased for clusters 1 and 2, or decreased for cluster 3 with decreasing pH.

**Figure 4.**
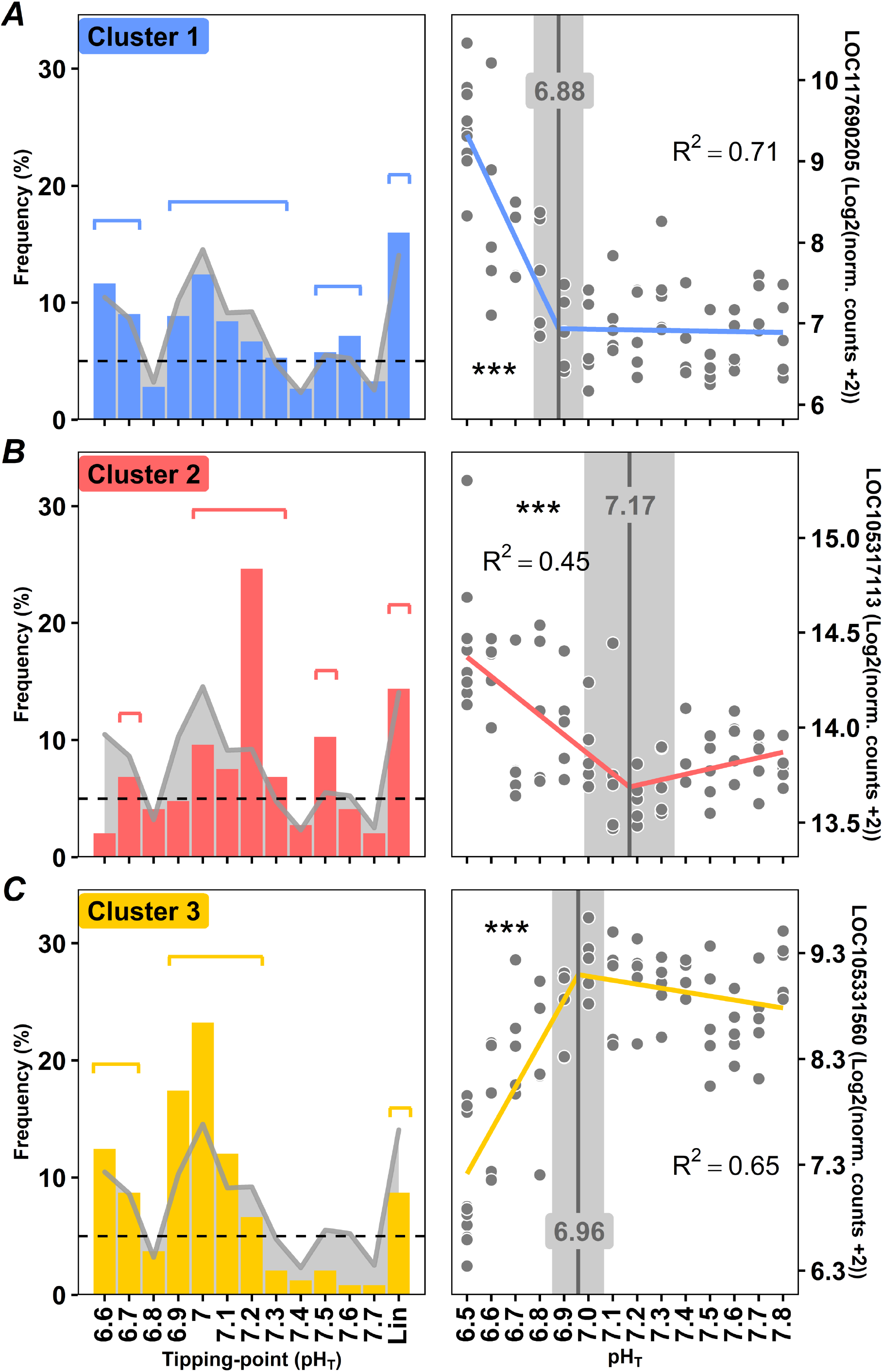
Tipping points of oyster transcriptome. (A-C) Frequency distribution of tipping-point for piecewise linear relationships (left side). Linear and loglinear models (no tipping-point) are under “Lin” name. Genes are grouped into three clusters of genes that co-vary together. The grey line indicates the distribution frequency of all genes irrespective of clusters. Groups of genes which exhibit neighboring tipping points with distribution frequencies > 5% (shown as a dotted line), were grouped together. The segments above the bars indicate the groups of genes on which GO analyses were conducted. In each case, the gene that best represents the cluster according to module membership, gene significance for pH and R^2^ is presented as a function of pH as an example (right side). Tipping points and their 95% confidence intervals are shown in grey. The significance levels of the slopes are presented using stars (P <0.001 ***, p<0.01 **, p<0.05 *). Gene names are: LOC117690205: monocarboxylate transporter 12-like, LOC105317113: 60S ribosomal protein L10a, LOC105331560: protocadherin Fat 4

Gene ontology analysis (GO) showed that most of the genes from cluster 1 (42%) exhibited tipping-points at pH 7.3-6.9 (Figure 4A) and were associated with GO terms corresponding to the regulation of RNA-transcription, cellular metabolism, macromolecule biosynthesis and negative regulation of cell-cell adhesion (Table 1). The expression of another group of genes from cluster 1 (14%) increased linearly (or loglinearly) with decreasing pH and was associated with GO terms corresponding to GTP-binding, GTPase activity and ribonucleotide metabolism (Table 1). The expression of most of the genes from cluster 2 (44%) exhibited tipping-points at pH 7.3-7.0, whereas another large group of genes (18%) increased linearly (or loglinearly) with decreasing pH (Figure 4B). These genes were associated with GO terms corresponding to ribosome synthesis, RNA-binding, translation and protein/amino acid synthesis (Table 1). The expression of 62% of the genes from cluster 3 exhibited tipping-points at pH 7.2-6.9 and decreased thereafter (Figure 4C). These genes were associated with GO terms corresponding to ion transport and more specifically to transmembrane cation transport (Table 1).

**Table 1.**
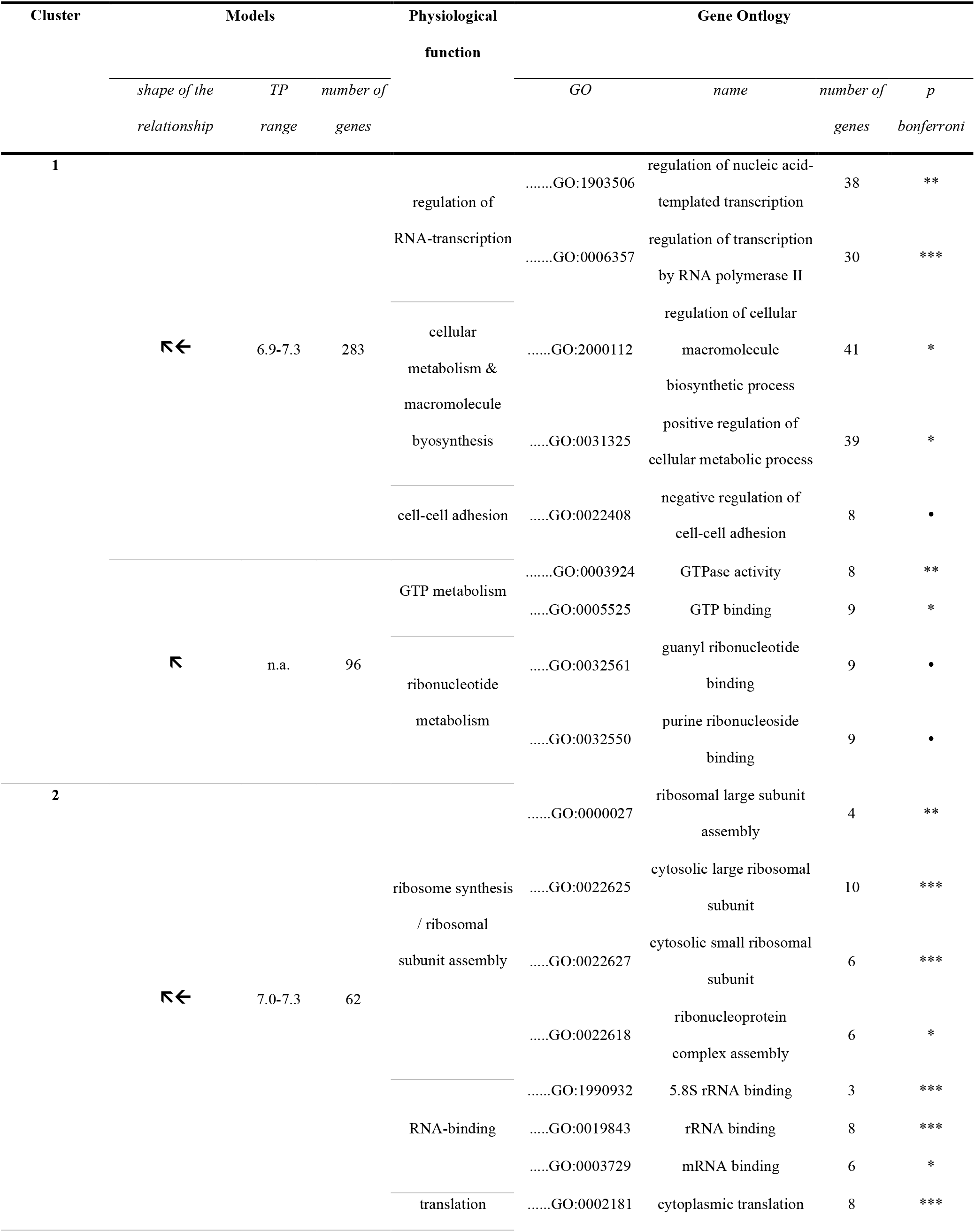
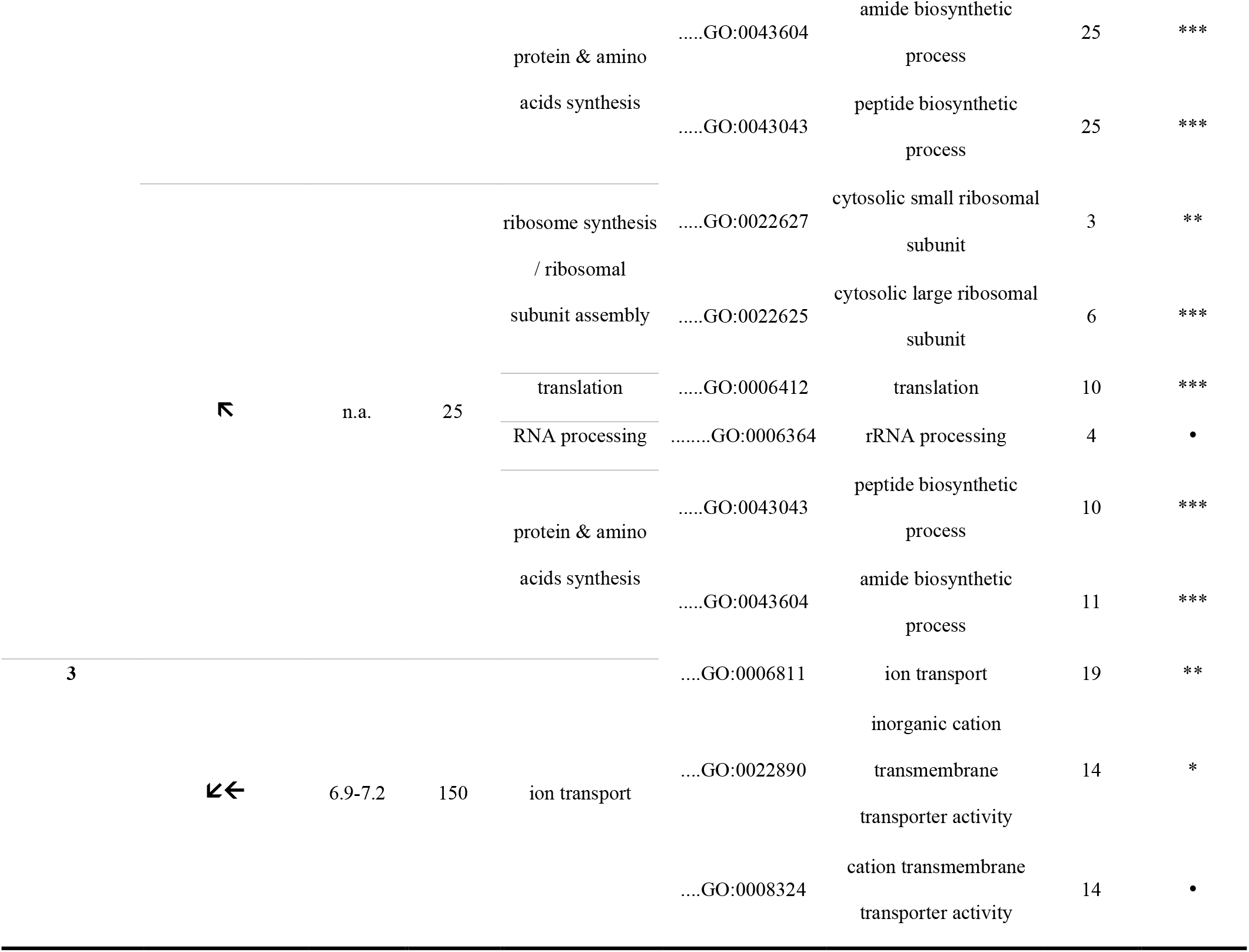
Gene ontology term enrichment and tipping point. Model characteristics, physiological function and gene ontology for each cluster. For each GO term, the P-value of Bonferroni test is displayed using symbols (P < 0.001 ***, < 0.01 **, < 0.05 *, < 0.1 .). Abbreviations: TP: tipping point

### Expression of genes related to biomineralization are affected by pH

We specifically examined the genes that are commonly reported in the scientific literature as being involved in the calcification process and in the formation of the organic matrix of the shell and periostracum. We found that the expression of 60% of the 38 genes involved in the calcification process that were retained by our models showed pH tipping points between 6.9 and 7.1 (Table 2). These genes encode for calcium-binding proteins, Ca^2+^ signaling pathway, amorphous calcium carbonate-binding proteins and ion transmembrane transporters. Most of these genes decreased below the tipping-point (n = 14) while others increased (n = 8). The expression of genes associated with the regulation of the synthesis of the shell organic matrix and the periostracum (n = 11) generally increased with decreasing pH (Table 2). The relationships were loglinear (n = 3) or piecewise (n = 8).

**Table 2.**
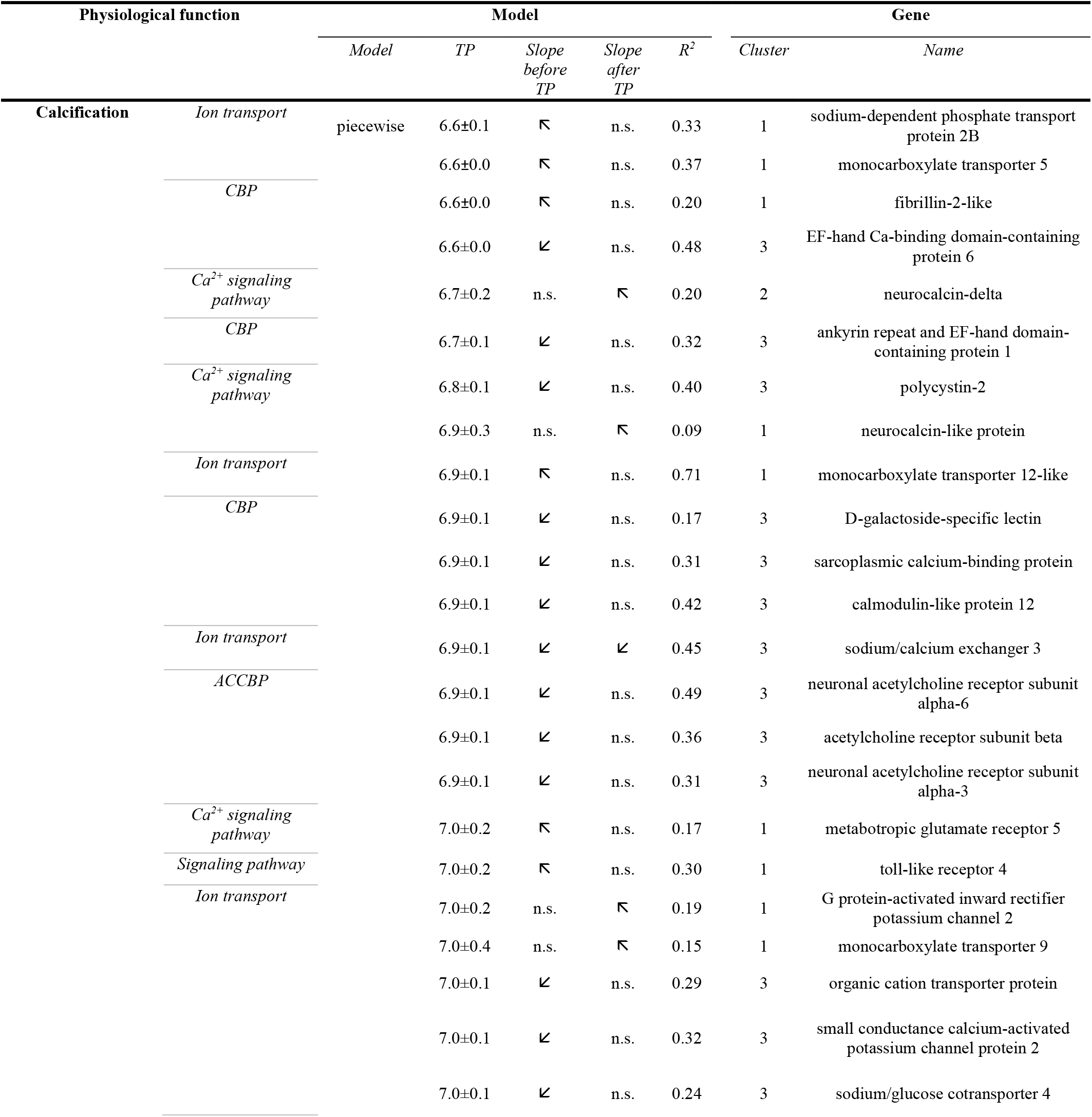
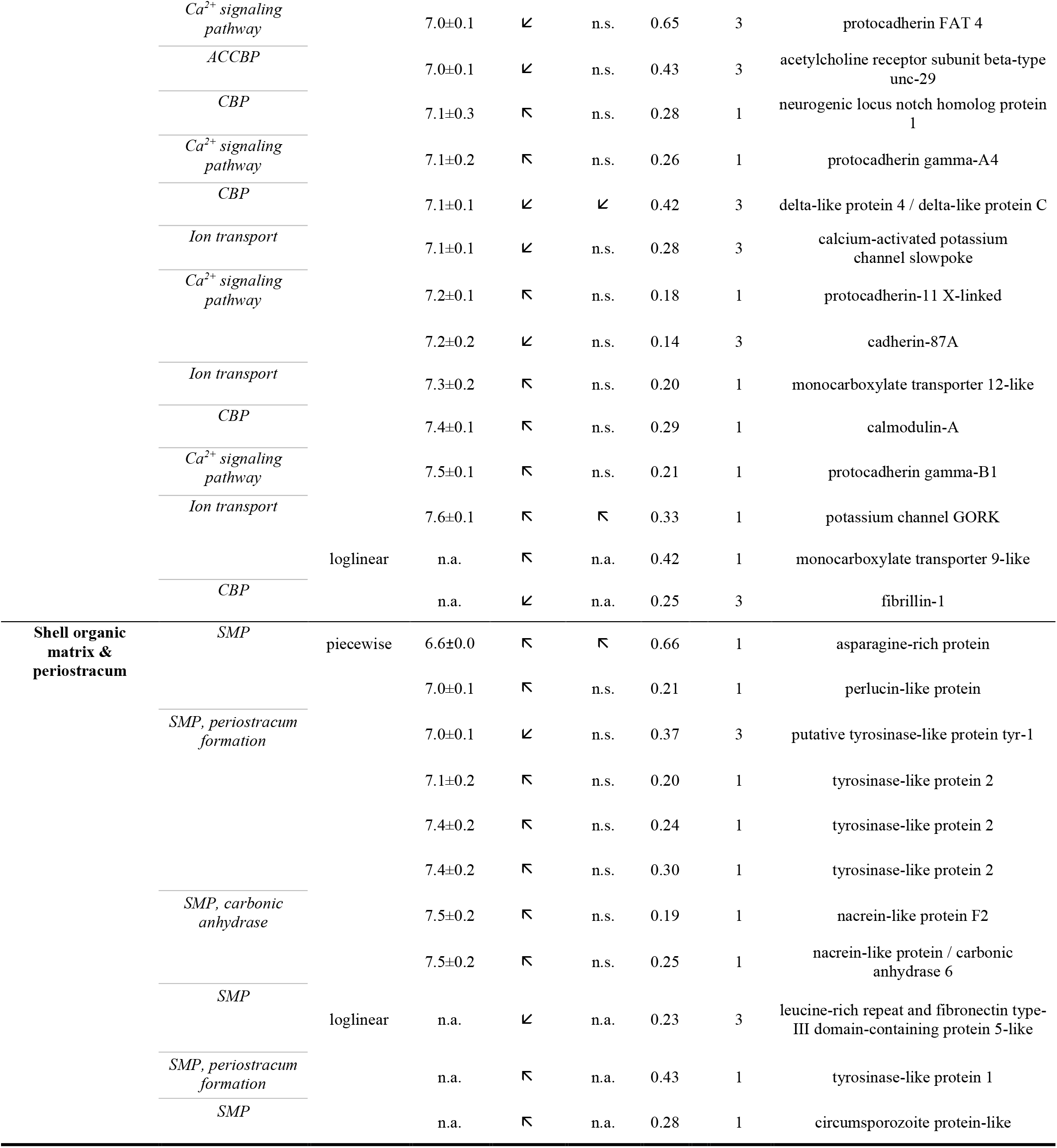
Tipping points of genes related to calcification and production of the shell organic matrix. Model characteristics and physiological function are reported for each gene. The slope inclination is displayed using arrows only if it is significant according to Student t tests (P < 0.05). Abbreviations: TP: tipping point, CBP: *calcium-binding proteins*, ACCBP: *amorphous calcium carbonate-binding proteins*, SMP: *shell matrix proteins*.

Among these 38 genes, the family of genes that are the most represented are coding for acetylcholine receptors, monocarboxylate transporters and tyrosinase synthesis (Table 2). The expression of four genes coding for acetylcholine receptors decreased below tipping points at pH 6.9-7.0. The expression of five genes coding for monocarboxylate transporters increased with decreasing pH, and four of them showed pH tipping points between 7.3 and 6.6. The expression of four genes associated to tyrosinase synthesis early increased with decreasing pH and three of them showed tipping points between pH 7.4 and 7.1.

## Discussion

Here we provide the first reaction norm of juvenile oyster *C. gigas* assessed by combining macro-physiological traits and micro-molecular characteristics. This novel approach applied to juvenile oysters exposed to a wide range of pH conditions revealed that macro-physiological parameters such as growth, calcification, food intake, and respiration exhibited pH tipping points that coincide with a major reshuffling in membrane lipids and transcriptome that was previously unappreciated, but others like shell parameters seemed uncoupled from global effect (Figure 5). This comprehensive view of the animal response to environment sheds new light on adaptation potential to climate change.

**Figure 5.**
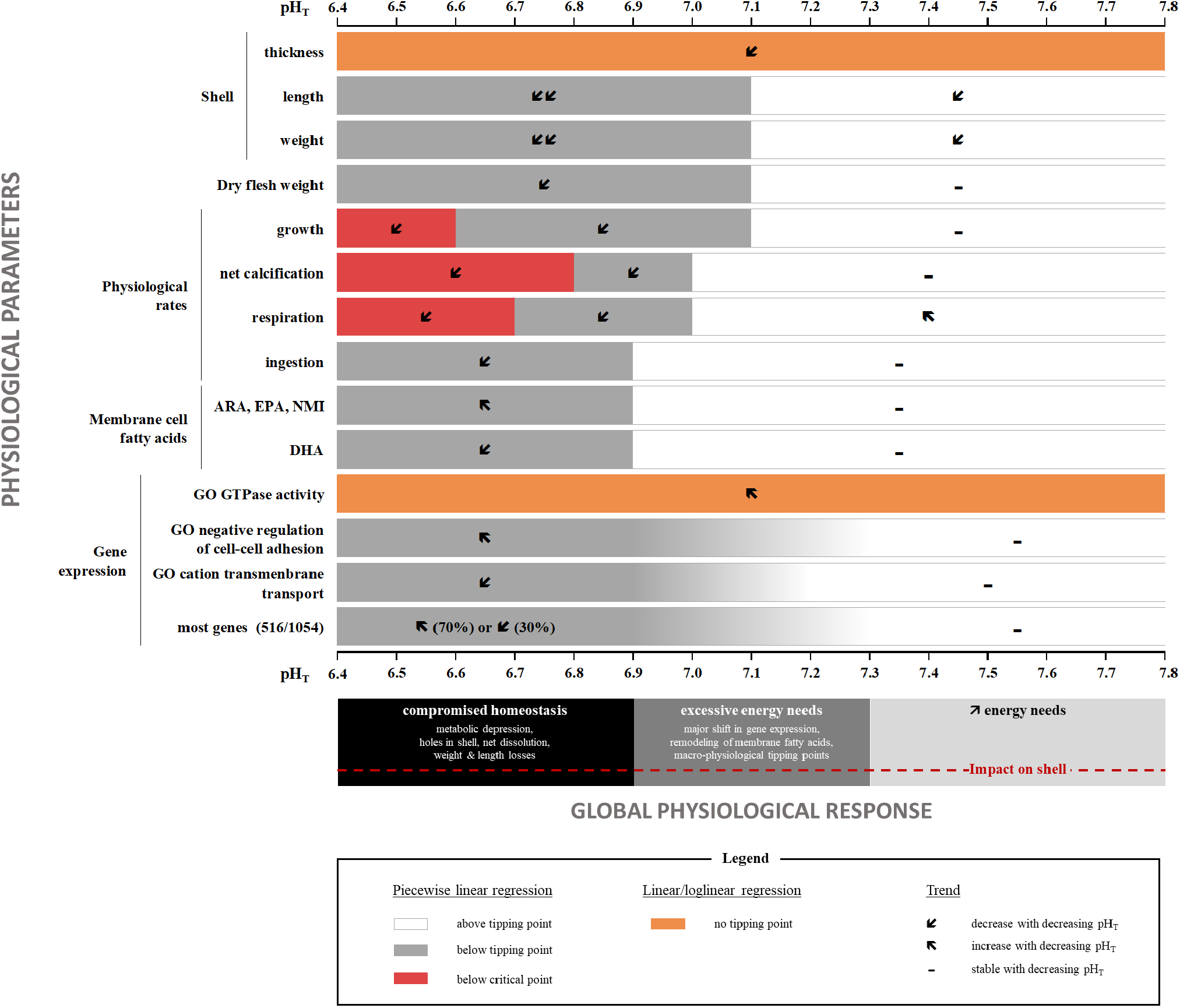
Graphical summary of the reaction norm of juvenile oysters *Crassostrea gigas* over a wide range of pH conditions. Abbreviations: GO, gene ontology term; ARA, arachidonic acid, EPA, ecosapentaenoic acid, DHA, docosahexaenoic acid, NMI, non-methylene interrupted.

We first identify a global tolerance threshold for juvenile *C. gigas* at pH 6.9-7.3, corresponding to the tipping points identified for the majority of the examined parameters. In line with this, several studies based on IPCC scenario assumed the existence of tipping points below pH 7.5-7.4 for growth, calcification and reproduction in oysters and mussels (Boulais et al., 2017; Fitzer et al., 2014; Lannig et al., 2010). Below tipping points, growth, calcification, respiration and feeding rate decreased markedly, likely reflecting increasing maintenance cost up until reaching critical points that are representative of metabolic depression (Michaelidis et al., 2005; Sokolova, 2021). However, these pH tipping points are well below IPCC projections for 2100 and have unlikely been encountered by mollusks since their appearance on earth 500 million years ago when CO_2_ levels were much higher (IPCC, 2019; Krissansen-Totton et al., 2018). Also, oyster populations do not generally experience such low pH conditions in their habitats (https://wwz.ifremer.fr/cocorico2_en/Donnees). Therefore, juvenile oysters appear to be able to cope with near-future OA.

We show here that shell parameters, i.e. thickness, weight and length, were altered as soon as pH decreased from ambient levels, as already reported for many shellfish species (Byrne & Fitzer, 2019). Shell thickness is generally related to shell strength and therefore with protection against predation and resistance to mechanical stress related to wave exposure and aquaculture processes. Thus, moderate acidification increases predation risk of young *C. gigas* through reduction of shell strength (Wright et al., 2018). We also find that respiration rate first increased when pH decreased, suggesting that metabolism was readily impacted. However, body condition (dry flesh weight) and energy reserves (TAG-ST and carbohydrate, Supplementary Figure 3) were not affected by pH, likely because oysters were fed *ad libitum* (Leung et al., 2019; Thomsen et al., 2013). Finally, we report here a linear increase in the expression level of genes related to GTPase activity and GTP binding, which are characteristics of stress response in *Crassostrea* spp. (Yan et al., 2018). Overall, these changes suggest that longer exposure to moderate acidification (above tipping point) could impair overall oyster fitness.

We further show that this reduction in shell parameters was not related to metabolic depression or to changes in net calcification or global gene expression. In contrast, a recent meta-analysis on *C. gigas* suggests that OA induces a reduction in shell thickness that is related to an overall transcriptional change inducing metabolic depression and alteration of calcification rate (Ducker & Falkenberg, 2020). Here we revisit these assumptions and find in agreement with other prior studies that no metabolic disruptions occurred while shell strength decreased at pH 7.3-7.7 (NBS scale) (Timmins-Schiffman et al., 2014; Wright et al., 2018).

Concomitantly, a delamination of the periostracum, the organic coat covering the shell was observed at pH levels above tipping point, probably altering shell protection. This was reflected by an increase shell bleaching and increased expression of four genes coding for tyrosinases-like protein. Tyrosinases are involved in periostracum synthesis and the increase in expression of their related genes is often considered as a mechanism to limit the damage to the periostracum and the corrosion of the shell in reaction to OA (Hüning et al., 2013). Delamination of the periostracum due to moderate OA was frequently associated with alteration of shell properties such as weakening and thinning (Alma et al., 2020; Auzoux-Bordenave et al., 2019; Bressan et al., 2014; Coleman et al., 2014; Peck et al., 2016, 2018; Zhao et al., 2020). Our results agree with the idea that OA is more a dissolution problem than a biomineralization problem (Rajan et al., 2021).

This investigation of tipping points at the micro-scale also provides new insights on the physiological response of oysters to OA. For the first time, we find a major remodeling of membrane lipids in response to OA and observed a tipping point at pH 6.9. This remodeling consisted of decreasing the long chain PUFA DHA (22:6n-3) that is essential for growth and survival (Knauer & Southgate, 1999; Langdon & Waldock, 1981), at the benefit of eicosanoid precursors (ARA, 20:4n-6 and EPA, 20:5n-3) involved in the stress response (Delaporte et al., 2003). Since membrane fatty acid composition influences the activity of transmembrane proteins involved in ion transport (Hazel & Williams, 1990; Hochachka & Somero, 2002), it could also have been related to regulation of acid-base homeostasis and perhaps calcification under OA.

In line with this, we find an overall decrease in expression of genes related to cation transmembrane transport below tipping points at pH 6.9-7.2 that can be related to alteration of calcification and acid-base homeostasis (Zhao et al., 2020). Concomitantly, we observe an overall decrease in the expression of genes coding for proteins involved in the regulation of calcification though calcium signaling pathway, calcium homeostasis, calmodulin signaling pathway and regulation of calcium carbonate crystal growth (Feng et al., 2017; Wang et al., 2017; Zhao et al., 2020).

In our analysis, we also identify families of genes that must play an important role in oyster response to OA. Among these genes, we identify four genes coding for acetylcholine receptors that are known for regulating and ordering the formation of the shell micro-structure (Feng et al., 2017). We also find five monocarboxylate transporters coding genes, members of a family of proteins that act as co-transporters of protons H^+^, that may be involved in the regulation of acid-base homeostasis and calcification process (Tresguerres et al., 2020; Wang et al., 2020). These proteins also co-transport monocarboxylates such as lactate or pyruvate and were reported to be involved in the stress response of oysters through regulation of metabolic pathways (Ertl et al., 2019).

In conclusion, we show that juvenile *C. gigas* have a broad tolerance to ocean acidification, exhibiting tipping points around pH 7.3-6.9 for most parameters (Figure 5). Nonetheless, we observe that shell parameters change as soon as pH drops, well before tipping points are reached, suggesting animal fitness is likely to be affected. This thus raises concerns about the future of natural and farmed oyster populations in a high-CO_2_ world. This new framework for identification of tolerance threshold in organisms represents a breakthrough in the field of global change research. It was made possible by (1) combining reaction norm assessment and thorough molecular and biochemical analyses of animal responses, and (2) developing a procedure to analyze and synthesize omics data measured over an environmental range. We believe that such an integrative and holistic approach could now be applied to other organisms and integrate intra-specific variation, life-stage and other stressors such as temperature, nutrition, pollutants or oxygen levels.

## Acknowledgments

We are grateful for K. Lugue, J. Veillet, C. Quéré, V. Le Roy and I. Queau for their participation to data recording and the Ifremer staff for their involvement in oyster and algae production. We thank J. Thebault and E. Dabass for their advices in shell analysis. We are also grateful for G. Le Moullac for project conception and funding. This work was funded by the Ocean Acidification Program of the French Foundation for Research on Biodiversity (FRB - www.fondationbiodiversite.fr) and the French Ministère de la transition écologique.

## Author contributions

ML, CDP, and FP designed and conducted the experiment. All authors analyzed the data. ML performed statistics. ML and FP wrote the first draft of the document and all authors contributed and accepted it. FP and JL obtained the funding. This work is part of the PhD thesis of ML.

## Data availability

RNA-seq data have been made available through the SRA database (BioProject accession number PRJNA735889). Other data analyzed during this study are included in this published article and its supplementary information files or available through the SEANOE database (https://doi.org/10.17882/83294). Complementary information is available from the corresponding authors on reasonable request.

