## Supplemental file for "Revisiting tolerance to ocean acidification: insights from a new framework combining physiological and molecular tipping points of Pacific oyster"

^1^ LEMAR UMR6539, CNRS/UBO/IRD/Ifremer, ZI pointe du diable, CS 10070, F-29280 Plouzané, France.

^2^LOV UMR7093, Sorbonne Université, CNRS, Laboratoire d'Océanographie de Villefranche, Villefranche-sur-Mer, France.

^3^EIO UMR24, UPF/IRD/ILM/Ifremer, Labex CORAIL, Unité RMPF, Centre Océanologique du Pacifique, Vairao – BP 49 Vairao, Tahiti, Polynésie française.

**Supplementary Note 1.**

Temperature, salinity and O_2_ saturation levels were stable across the experimental period and similar among pH conditions (Supplementary Table 1). Seawater was undersaturated with respect to calcite and aragonite when pH was below 7.3 and 7.6, respectively (Supplementary Table 1). However, parameters of carbonate chemistry were sometimes similar for different pH conditions. This reflects the difference in the frequency of measurements between the parameters (twice daily for pH and weekly for TA). Also, TA decreased slightly but linearly with increasing pH (R^2^ = 0.82, slope = -0.08, P < 0.001) suggesting that seawater renewal was likely not sufficient to counteract both the uptake and the release of carbonate ions from net calcification at high pH and net dissolution at low pH, respectively (Supplementary Figure 1).

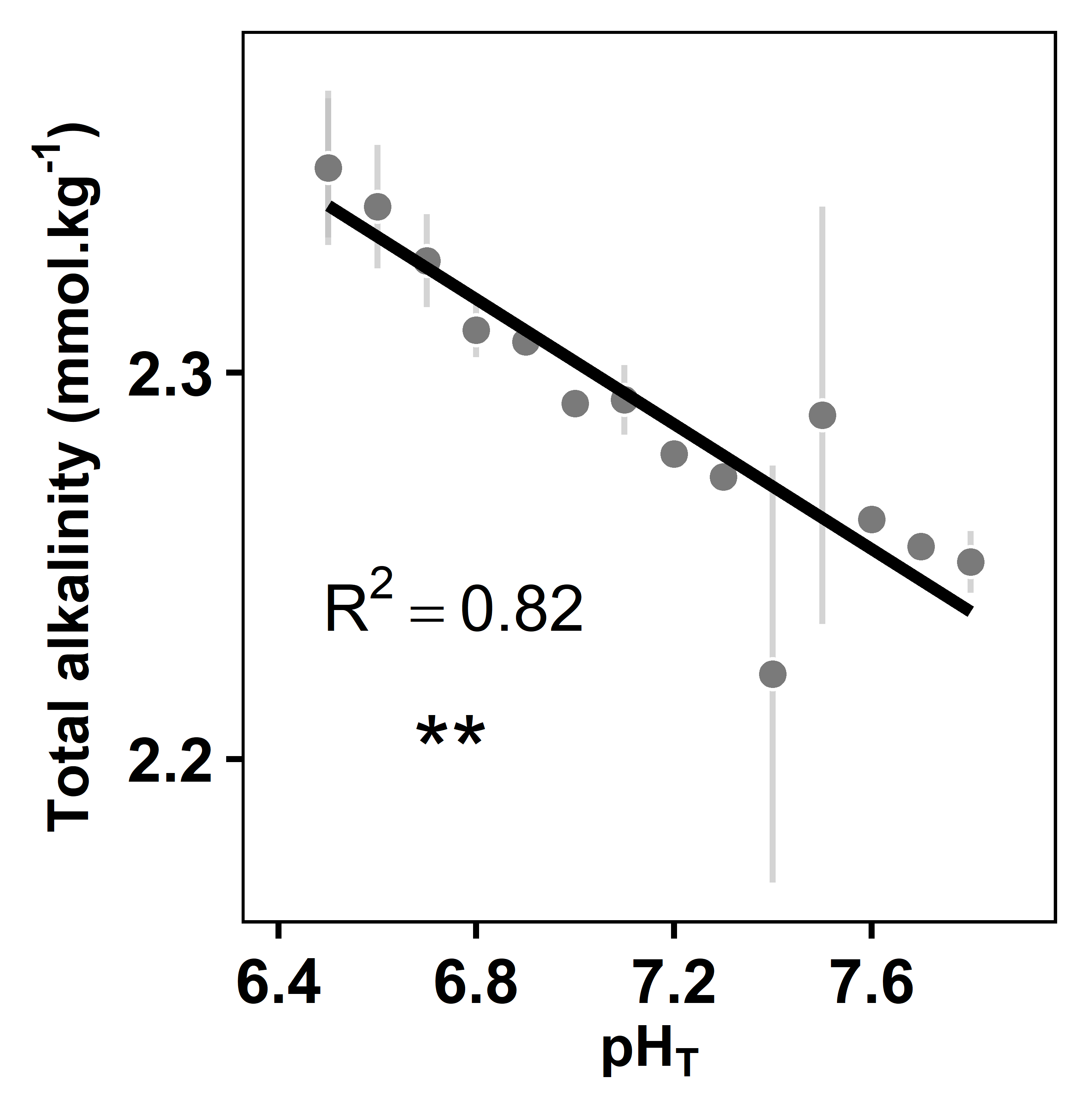

**Supplementary Figure 1.** Total alkalinity as a function of pH (total scale). Data are means ± SD values recorded during the 23 d of exposure. The significance level of the slope is presented using stars (P <0.001 ***, <0.01 **, <0.05 *, <0.1 **^.^**).

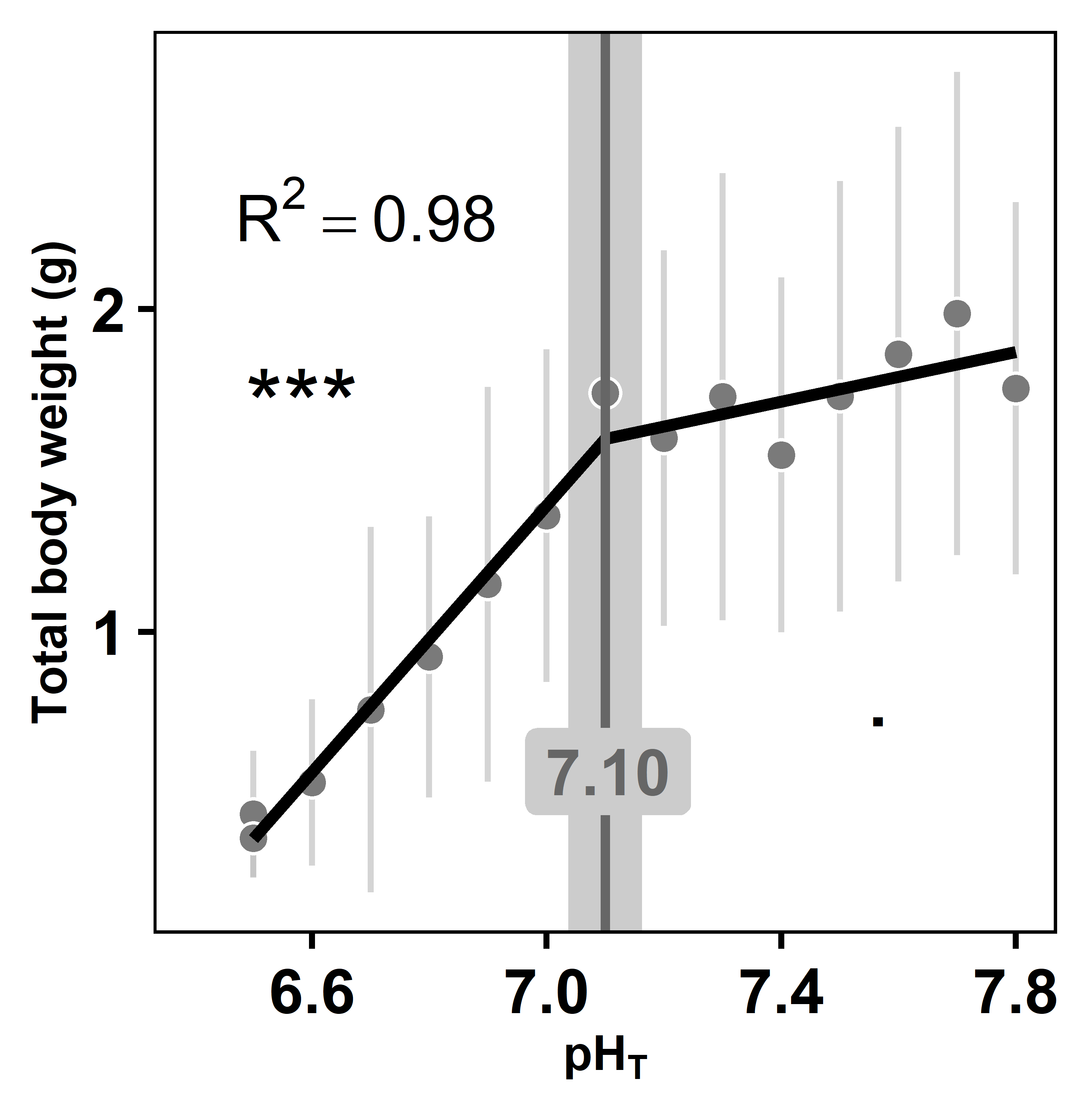

**Supplementary Figure 2.** Total body weight of oysters as a function of pH (pH on the total scale). Data are means ± SD. Tipping-point and its confidence intervals is shown in grey. The significance levels of the slopes are presented using stars (P < 0.001 ***, < 0.01 **, < 0.05 *, < 0.1 **^.^**).

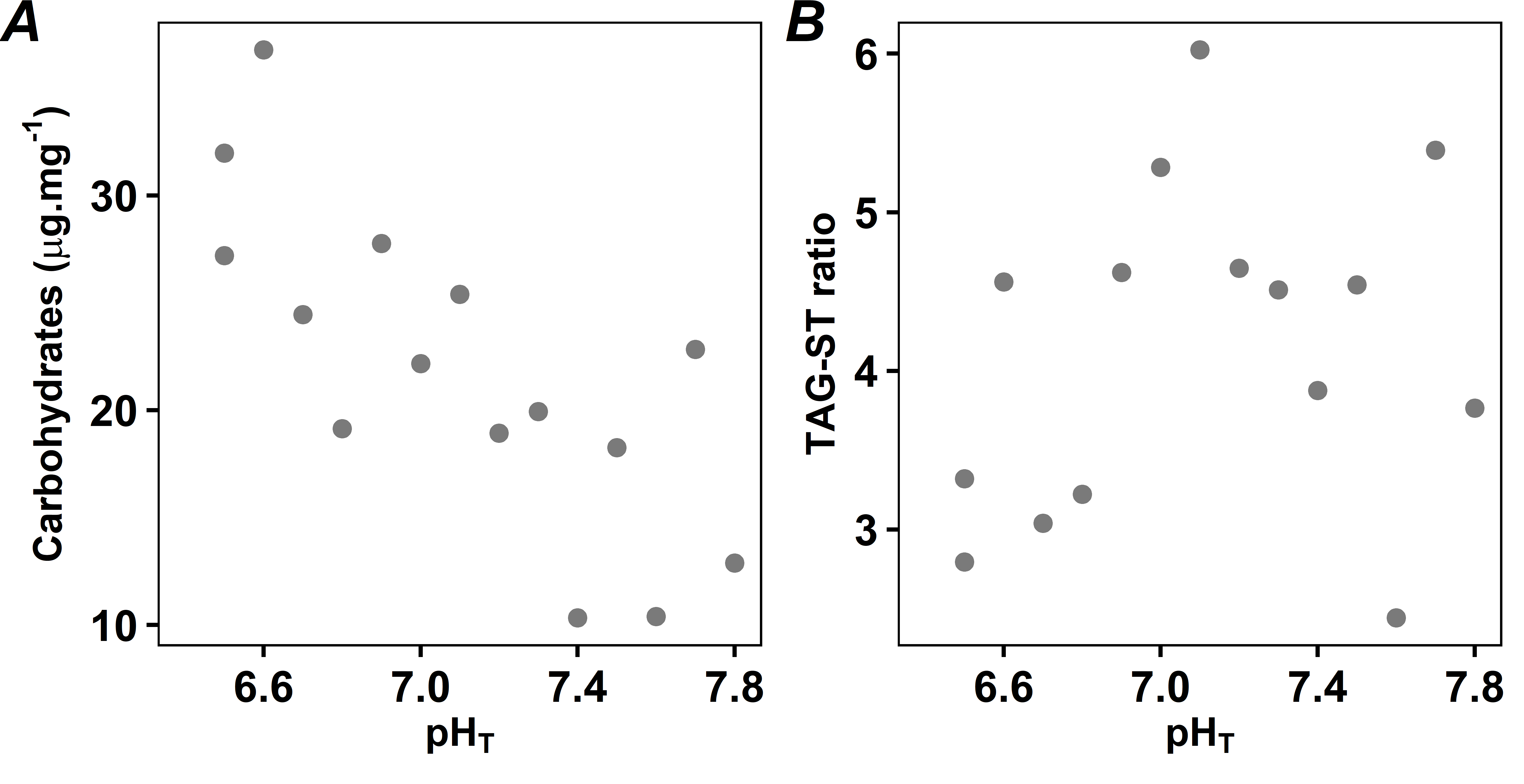

**Supplementary Figure 3.** Energy reserves of oysters as a function of pH (total scale). (A) total carbohydrates and (B) triacylglycerol/sterol ratio (TAG-ST). Relationships were not significant (P > 0.05).
**Supplementary Table 1** Seawater parameters measured during the experiment in each experimental tank, including three blanks without oysters. pH on the total scale (pH_T_), total alkalinity (TA), oxygen saturation (O_2_), salinity (S), temperature (T), calcite saturation state (Ω_CA_), aragonite saturation state (Ω_AR_), CO_2_ partial pressure (pCO_2_) and dissolved inorganic carbon (DIC) concentrations are shown (mean ± SD). pH, O_2_, S and T were measured twice daily and TA was measured once a week.

| *Nominal* |  | *Measured* | | | | |  | *Calculated* | | | |
| --- | --- | --- | --- | --- | --- | --- | --- | --- | --- | --- | --- |
| **pH_T_** |  | **pH_T_** | **TA** (μmol.kg^-1^) | **O_2_** (%sat) | **S** | **T (°C)** |  | **Ω_CA_** | **Ω_AR_** | **pCO_2_** (μatm) | **DIC** (μmol.kg^-1^) |
| Blank |  | 8.0±0.1 | 2331±7 | 101.6±0.5 | 35.7±0.1 | 21.8±0.1 |  | 3.9±0.4 | 2.5±0.3 | 552±82 | 2111±31 |
| Blank |  | 8.0±0.1 | 2328±3 | 101.5±0.4 | 35.7±0.1 | 21.7±0.1 |  | 3.9±0.4 | 2.5±0.3 | 551±82 | 2108±29 |
| Blank |  | 7.9±0.2 | 2331±6 | 101.4±0.4 | 35.7±0.1 | 21.7±0.1 |  | 3.9±0.4 | 2.5±0.3 | 552±82 | 2111±29 |
| 7.8 |  | 7.8±0.1 | 2251±8 | 97.6±1.8 | 35.7±0.1 | 21.8±0.1 |  | 3.1±0.3 | 2.0±0.2 | 695±103 | 2082±33 |
| 7.7 |  | 7.7±0.1 | 2255±2 | 97.6±1.6 | 35.7±0.1 | 21.8±0.1 |  | 2.2±0.3 | 1.4±0.2 | 1070±161 | 2152±19 |
| 7.6 |  | 7.6±0.1 | 2262±2 | 96.7±2.6 | 35.7±0.1 | 21.7±0.1 |  | 1.5±0.0 | 1.0±0.0 | 1617±1 | 2217±2 |
| 7.5 |  | 7.5±0.1 | 2289±54 | 96.5±2.2 | 35.7±0.1 | 21.7±0.1 |  | 1.3±0.0 | 0.8±0.0 | 2091±49 | 2276±55 |
| 7.4 |  | 7.4±0.1 | 2222±54 | 97.1±2.1 | 35.7±0.1 | 21.7±0.1 |  | 1.1±0.1 | 0.7±0.1 | 2215±352 | 2219±66 |
| 7.3 |  | 7.3±0.0 | 2273±3 | 96.7±2.3 | 35.7±0.1 | 21.7±0.1 |  | 1.0±0.0 | 0.7±0.0 | 2644±4 | 2292±3 |
| 7.2 |  | 7.2±0.1 | 2279±3 | 96.8±2.2 | 35.7±0.1 | 21.8±0.1 |  | 0.6±0.0 | 0.4±0.0 | 4276±7 | 2371±3 |
| 7.1 |  | 7.1±0.1 | 2293±9 | 96.8±1.7 | 35.7±0.1 | 21.7±0.1 |  | 0.6±0.1 | 0.4±0.0 | 5067±671 | 2414±30 |
| 7.0 |  | 7.0±0.0 | 2292±3 | 97.2±1.5 | 35.7±0.1 | 21.7±0.1 |  | 0.6±0.1 | 0.4±0.0 | 5062±664 | 2413±27 |
| 6.9 |  | 6.9±0.1 | 2308±2 | 97.7±1.6 | 35.7±0.1 | 21.7±0.1 |  | 0.4±0.0 | 0.2±0.0 | 8160±1063 | 2536±38 |
| 6.8 |  | 6.8±0.1 | 2311±7 | 97.6±1.9 | 35.7±0.1 | 21.7±0.2 |  | 0.3±0.0 | 0.2±0.0 | 8777±26 | 2560±8 |
| 6.7 |  | 6.7±0.1 | 2329±12 | 98.2±1.3 | 35.7±0.1 | 21.7±0.1 |  | 0.3±0.0 | 0.2±0.0 | 11172±60 | 2656±13 |
| 6.6 |  | 6.6±0.1 | 2343±16 | 98.9±1.0 | 35.7±0.1 | 21.7±0.2 |  | 0.2±0.1 | 0.1±0.0 | 15691±3912 | 2813±135 |
| 6.5 |  | 6.5±0.2 | 2353±20 | 99.2±0.8 | 35.7±0.1 | 21.7±0.2 |  | 0.2±0.0 | 0.1±0.0 | 19865±4915 | 2953±164 |
| 6.4 |  | 6.5±0.3 | 2353±18 | 99.2±0.9 | 35.7±0.1 | 21.7±0.2 |  | 0.1±0.0 | 0.1±0.0 | 23079±5471 | 3054±183 |

**Supplementary Table 2** Relative contribution of fatty acids to membrane. Their contribution (%) and Pearson’s correlation to the first principal component (PC1) of the PCA are showed.

| **Average pH_T_** | |  | **Fatty acid (% total fatty acids)** | | | |  |  |  |  |  |  |
| --- | --- | --- | --- | --- | --- | --- | --- | --- | --- | --- | --- | --- |
|  |  |  | **14:0** | **16:0** | **16:1n-7** | **18:0 dma** | **18:0** | **18:1n-9** | **18:1n-7** | **18:2n-6** | **18:3n-3** | **18:4n-3** |
| 7.8 | |  | 2.7 | 14.1 | 1.7 | 9.7 | 5.6 | 1.8 | 5.3 | 2.2 | 1.3 | 1.7 |
| 7.7 | |  | 2.7 | 12.5 | 2.1 | 9.2 | 3.6 | 1.9 | 6.0 | 2.3 | 1.4 | 1.8 |
| 7.6 | |  | 2.5 | 13.7 | 1.6 | 9.8 | 4.0 | 1.9 | 5.4 | 2.4 | 1.3 | 1.6 |
| 7.5 | |  | 2.0 | 12.5 | 1.6 | 10.8 | 4.4 | 1.6 | 5.6 | 1.8 | 1.2 | 1.4 |
| 7.4 | |  | 2.2 | 13.0 | 1.7 | 10.8 | 4.7 | 1.6 | 5.5 | 1.9 | 1.3 | 1.5 |
| 7.3 | |  | 2.2 | 13.7 | 1.6 | 9.4 | 4.3 | 1.8 | 5.1 | 2.2 | 1.3 | 1.6 |
| 7.2 | |  | 2.5 | 12.8 | 1.8 | 9.9 | 3.9 | 1.7 | 5.5 | 2.2 | 1.4 | 1.9 |
| 7.1 | |  | 2.3 | 12.0 | 1.8 | 11.2 | 4.6 | 1.7 | 5.2 | 1.9 | 1.4 | 1.7 |
| 7.0 | |  | 2.4 | 12.7 | 1.8 | 10.7 | 4.3 | 1.6 | 4.4 | 1.9 | 1.2 | 1.6 |
| 6.9 | |  | 2.0 | 11.6 | 1.7 | 11.3 | 4.8 | 1.8 | 6.0 | 2.0 | 1.3 | 1.6 |
| 6.8 | |  | 2.0 | 12.0 | 1.7 | 9.8 | 4.7 | 1.8 | 5.9 | 2.0 | 1.1 | 1.4 |
| 6.7 | |  | 1.5 | 11.9 | 1.7 | 10.6 | 6.3 | 1.7 | 6.2 | 1.6 | 0.8 | 1.0 |
| 6.6 | |  | 2.0 | 11.8 | 1.9 | 9.3 | 4.6 | 2.0 | 6.7 | 2.1 | 1.0 | 1.4 |
| 6.5 | |  | 1.8 | 10.8 | 2.3 | 10.0 | 2.4 | 1.7 | 6.7 | 1.4 | 0.7 | 1.0 |
| 6.5 | |  | 1.5 | 11.3 | 2.0 | 9.0 | 5.7 | 1.7 | 6.5 | 1.3 | 0.7 | 0.7 |
| **PCA** | Contribution to PC1 (%) |  | 1.9 | 13.6 | 0.5 | 0.7 | 0.0 | 0.0 | 4.6 | 1.7 | 1.1 | 1.8 |
|  | Correlation to PC1 (Pearson) |  | 0.8 | 0.9 | -0.8 | 0.2 | 0.0 | 0.0 | -0.7 | 0.8 | 0.9 | 0.9 |
| **Average pH_T_** | |  | **Fatty acid (% total fatty acids)** | | | |  |  |  |  |  |  |
|  |  |  | **20:1 dma** | **20:0** | **20:1n-11** | **20:1n-7** | **20:4n-6** | **20:5n-3** | **22:2 NMI_i,j_** | **22:5n-6** | **22:6n-3** |  |
| 7.8 | |  | 1.9 | 1.1 | 1.7 | 3.8 | 4.5 | 8.7 | 5.0 | 3.2 | 15.8 |  |
| 7.7 | |  | 1.9 | 1.1 | 1.9 | 3.7 | 4.9 | 9.9 | 5.0 | 3.0 | 15.2 |  |
| 7.6 | |  | 1.9 | 1.2 | 1.7 | 4.1 | 4.4 | 9.5 | 4.5 | 3.0 | 16.3 |  |
| 7.5 | |  | 2.1 | 1.1 | 2.0 | 4.0 | 4.5 | 9.6 | 5.4 | 3.1 | 16.1 |  |
| 7.4 | |  | 2.1 | 1.1 | 2.0 | 4.1 | 4.4 | 9.2 | 5.4 | 3.2 | 16.6 |  |
| 7.3 | |  | 1.8 | 1.2 | 1.8 | 3.9 | 4.7 | 10.1 | 4.7 | 2.9 | 16.0 |  |
| 7.2 | |  | 1.9 | 1.1 | 2.0 | 4.1 | 4.5 | 9.6 | 5.1 | 3.0 | 16.3 |  |
| 7.1 | |  | 2.1 | 1.2 | 2.2 | 4.2 | 4.6 | 9.6 | 6.0 | 2.8 | 14.0 |  |
| 7.0 | |  | 2.1 | 1.1 | 2.1 | 4.3 | 4.9 | 9.9 | 5.7 | 3.0 | 15.7 |  |
| 6.9 | |  | 1.9 | 1.1 | 2.1 | 4.0 | 4.8 | 9.7 | 5.4 | 3.1 | 15.5 |  |
| 6.8 | |  | 2.0 | 1.2 | 2.0 | 4.4 | 5.1 | 9.9 | 5.7 | 3.0 | 14.7 |  |
| 6.7 | |  | 2.3 | 1.2 | 2.1 | 4.5 | 5.6 | 10.3 | 5.9 | 2.5 | 13.7 |  |
| 6.6 | |  | 1.9 | 1.4 | 1.9 | 4.3 | 5.4 | 9.7 | 5.9 | 2.7 | 14.3 |  |
| 6.5 | |  | 2.8 | 1.2 | 2.2 | 4.9 | 6.5 | 11.1 | 6.6 | 2.0 | 12.5 |  |
| 6.5 | |  | 2.5 | 1.4 | 2.1 | 5.4 | 6.9 | 11.1 | 6.4 | 2.0 | 11.8 |  |
| **PCA** | Contribution to PC1 (%) |  | 1.2 | 0.1 | 0.2 | 3.2 | 11.8 | 6.1 | 5.8 | 2.8 | 43.0 |  |
|  | Correlation to PC1 (Pearson) |  | -0.8 | -0.7 | -0.6 | -0.9 | -1.0 | -0.9 | -0.9 | 0.9 | 1.0 |  |

**Supplementary Table 3** Number of genes modelled against pH. Each step of filtering and the corresponding number of selected genes are specified.

| **WGCNA cluster of gene** | | **correlated with pH** | **modelled & slope ≠ 0** | **model** | |
| --- | --- | --- | --- | --- | --- |
| 1 | 3380 | 1691 | 664 | piecewise | 567 |
|  |  |  |  | linear | 97 |
| 2 | 723 | 343 | 148 | piecewise | 130 |
|  |  |  |  | linear | 18 |
| 3 | 2073 | 623 | 242 | piecewise | 221 |
|  |  |  |  | linear | 21 |

**Supplementary Table 4** Seawater parameters during the incubations. pH on the total scale (pH_T_), calcite saturation state (Ω_CA_), total alkalinity (TA), oxygen saturation (O_2_), salinity (S) and temperature (T). Average represents the mean values between the onset and the end of the incubations. Δ represents the difference between the end and the onset of the incubations. Values are mean ± SD.

| **Nominal pH_T_** | **Average pH_T_** | **Δ pH_T_** | **Average Ω_CA_** | **Δ Ω_CA_** | **Δ TA** (μmol.kg^-1^) | **Δ O_2_** (%sat) | **Δ NH_4_^+^** (µmol.L^-1^) | **Average S** | **Average T** (°C) |
| --- | --- | --- | --- | --- | --- | --- | --- | --- | --- |
| Blank | 8.0±0.0 | 0.0 | 4.4±0.0 | 0.0 | -3 | 2 | 0.0 | 35.7±0.0 | 21.7±0.1 |
| Blank | 8.0±0.0 | 0.0 | 4.3±0.0 | 0.0 | 1 | 1.7 | -0.1 | 35.7±0.0 | 21.6±0.1 |
| Blank | 8.0±0.0 | 0.1 | 4.4±0.0 | 0.0 | 0 | 1.9 | 0.3 | 35.7±0.0 | 21.7±0.1 |
| 7.8 | 7.9±0.1 | 0.0 | 3.1±0.5 | 0.7 | 80 | 3 | -1.1 | 35.7±0.0 | 21.7±0.1 |
| 7.7 | 7.6±0.1 | 0.1 | 1.7±0.3 | 0.4 | 65 | 6.5 | -1.2 | 35.7±0.0 | 21.6±0.1 |
| 7.6 | 7.5±0.1 | 0.0 | 1.4±0.2 | 0.3 | 60 | 5 | -1.0 | 35.7±0.0 | 21.7±0.1 |
| 7.5 | 7.4±0.1 | -0.1 | 1.1±0.2 | 0.3 | 72 | 7.9 | 1.4 | 35.7±0.0 | 21.7±0.1 |
| 7.4 | 7.3±0.1 | 0.1 | 0.9±0.2 | 0.2 | 72 | 7.3 |  | 35.7±0.0 | 21.5±0.1 |
| 7.3 | 7.3±0.0 | -0.2 | 1.0±0.0 | 0.0 | 61 | 6.1 | -1.0 | 35.7±0.0 | 21.7±0.1 |
| 7.2 | 7.1±0.0 | 0.0 | 0.6±0.0 | 0.0 | 38 | 6.7 | -0.8 | 35.7±0.0 | 21.7±0.1 |
| 7.1 | 7.1±0.0 | 0.0 | 0.6±0.0 | 0.0 | 44 | 5.5 | -1.1 | 35.7±0.0 | 21.6±0.1 |
| 7.0 | 7.0±0.0 | 0.0 | 0.5±0.0 | 0.0 | 36 | 6.1 | -2.0 | 35.7±0.0 | 21.7±0.1 |
| 6.9 | 6.9±0.0 | 0.1 | 0.4±0.0 | 0.0 | 19 | 5.5 | -0.9 | 35.7±0.0 | 21.7±0.1 |
| 6.8 | 6.9±0.1 | -0.1 | 0.4±0.1 | -0.1 | 2 | 4.9 | -1.0 | 35.7±0.0 | 21.6±0.1 |
| 6.7 | 6.8±0.1 | -0.1 | 0.3±0.0 | -0.1 | 13 | 3.1 | -1.1 | 35.7±0.0 | 21.7±0.1 |
| 6.6 | 6.8±0.1 | -0.2 | 0.3±0.1 | -0.2 | 0 | 2.7 | -0.6 | 35.7±0.0 | 21.6±0.1 |
| 6.5 | 6.7±0.1 | 0.1 | 0.3±0.1 | -0.1 | -12 | 2.4 | -0.6 | 35.7±0.0 | 21.7±0.1 |
| 6.4 | 6.6±0.1 | 0.0 | 0.2±0.0 | 0.0 | -18 | 1.5 | -0.6 | 35.7±0.0 | 21.6±0.1 |
